# CView: A network based tool for enhanced alignment visualization

**DOI:** 10.1101/2022.01.17.476623

**Authors:** Raquel Linheiro, Diana Lobo, Stephen Sabatino, John Archer

## Abstract

To date basic visualization of sequence alignments have largely focused on displaying per-site columns of nucleotide, or amino acid, residues along with associated frequency summarizations. The persistence of this tendency to the more recent tools designed for the viewing of mapped read data indicates that such a perspective not only provides a reliable visualization of per-site alterations, but also offers implicit reassurance to the end user in relation to data accessibility. However, the initial insight gained is limited, something that is especially true when viewing alignments consisting of many sequences representing differing factors, such as geographical location, date and subtype. A basic alignment viewer can have potential to increase initial insight through visual enhancement, whilst not delving into the realms of complex sequence analysis. Here we present CView, a visualizer that expands on the per-site representation of residues through the incorporation of a dynamic network that is based on the summarization of diversity present across different regions of the alignment. Within the network nodes are based on the clustering of sequence fragments spanning windows that are placed consecutively along the alignment. Edges are placed between nodes of neighbouring windows where they share sequence id’s. Thus, if a single node is selected on the network, then the relationship that all sequences passing through that node have to other regions of diversity within the alignment can be instantly observed through the tracing of paths. In addition to augmenting visual insight, CView provides many export features including variant summarization, per-site residue and kmer frequency matrixes, consensus sequence generation, alignment dissection as well as general sequence clustering, each of which are useful across a range of research areas. The software has been designed to be user friendly, intuitive and interactive. It, along with source code, a quick start guide and test data, are available through the SourceForge project page: https://sourceforge.net/projects/cview/.

## Introduction

Tools developed to visualize local sets of aligned sequences, such as those produced by multiple sequence aligners including MUSCLE [1] and Clustal W [2], have focused largely on displaying columns of nucleotide or amino acid characters, and highlighting the differences between such characters primarily through the use of colour [3–7]. More advanced sequence management and analysis packages, such as Geneious [8] and Mega [9], as well as the more recent tools designed for basic visualization of mapped read data, including IGV [10], GenomeView [11] and Tablet [12], incorporate a wide array of analysis, summarization and annotation options, but in terms of basic visualization they follow a similar approach. It is evident that direct observation of residues, as well as the general per-site based summarization, not only provides an accessible view of per-site alterations between sequences within the alignment but also at times gives the end user a level of reassurance in relation to their data. However, initial insight gained in relation to the overall alignment is limited, especially when viewing alignments consisting of many sequences representing varying factors of interest such as geographical location, subtype, treatment strategy, compartment and date / time-point. Basic alignment visualization should have the potential to increase the level of initial insight within sequence datasets whilst not delving into the realms of more complex sequence analysis. Here we present CView, a simple multiple sequence alignment visualizer that incorporates a dynamic network that is based on a summarization of the diversity across different regions of the alignment. The immediate coupling of aligned sequences to such a network provides a way of visually tracking the context of observed diversity within characters that are currently onscreen to that of the surrounding regions of the alignment not currently in view. This provides the user with an increased intuitive and visual summarization of the context of this diversity.

CView provides a range of export features that can be applied to the entire alignment, to a specified region of the alignment, or to a specified region in conjunction with a specified subset of sequences. Such export features include: variant summarization, per-site character and kmer frequency matrixes, clustered sequences, pairwise-distance matrixes as well as consensus sequence generation. For example, when the variant summarization option is selected a list of variants spanning the user-specified region, within the user-specified group of sequences, is created by identifying all unique forms and associating each with their frequency of occurrence. A list of the original sequence titles represented by each variant is also maintained. Such a feature has use in the tracking of viral populations, for example in searching for the presence of genotypic alterations such as those associated with immune escape [13], drug resistance [14], or co-receptor usage [15,16]. Additionally, this feature has use in both clinical [17], and environmental metagenomics [18–20], where the summarization of populations of microbes is of interest. Each such export option is described detail within the user manual that is available on the SourceForge wiki located at https://sourceforge.net/p/cview/wiki/Help/. Aside from features related to the extraction of secondary information from the alignment, CView provides the ability to dissection the alignment into subsets of sequences and regions; a task that is often laborious in the absence of a background in script development. For example, a user can export a specified region of sequences associated with a specific time-point, geographical location or body compartment, as long as the sequence titles have been labelled with such information. Such labelling is often as standard output feature of many sequence repositories, for example from the Los Alamos HIV sequence database the user can select options such as subtype, patient code, country and year to be included within the title of each sequence [21], but such information may also be part of experimental design such as compartment [22] or time-point [23].

Within CView the associated network is displayed directly below the alignment. This network is based on the clustering of sequences within windows placed consecutively across the alignment, where each cluster becomes a node. Edges are placed between nodes of neighbouring windows where they share differing regions of the same sequence(s). Thus, if a single node within a diverse region of the alignment is selected, then the relationship that all sequences passing through that node have to other regions of diversity within the alignment can be instantly observed. Here we describe how these networks are constructed and graphically displayed. The clustering threshold used during network construction, as well as the number and width of windows, are specified on the user-interface through a series of user-friendly slider bars. Alterations are updated in real time, which allows the user to rapidly explore the visualization of variable regions across the alignment. The software has been designed to be user friendly, intuitive and interactive and it, along with source code, a quick start guide and test data, is available through the SourceForge project page: https://sourceforge.net/projects/cview/.

## Methods

### Implementation

The interface has been designed for simplicity and clarity. It consists of four basic areas of user-interaction (figure 1) which are: (1) sequence view, (2) network view, (3) navigation and control and, (4) menu driven outputs.

**Figure 1:**
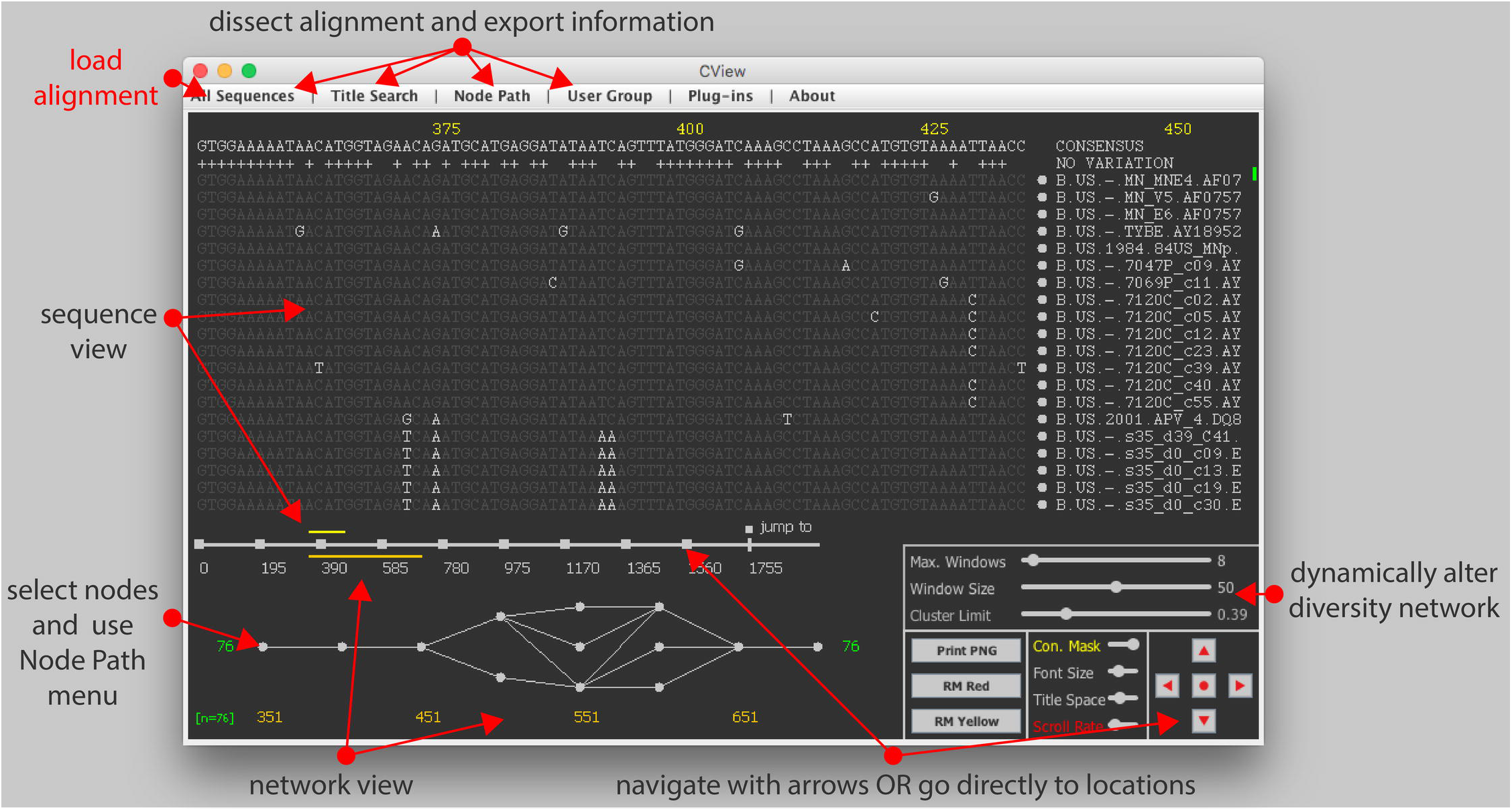
CView interface. The four main areas of the CView interface are depicted. These are sequence view, network view, control panel and the top menu. The yellow numbers on the top indicate the sites of the alignment that are currently in view. These correspond to the yellow bar on the top of the location indicator. The orange numbers along the bottom indicate the locations of the windows that nodes within the network are dependent on. These window locations correspond to the area that the orange bar located the under the location indicator covers. Grey dots indicate (selectable) nodes within windows. The squares along the location indicator can also be selected in order to jump directly to the indicated co-ordinates. The red text around the outside of the interface describes the main features.

#### (1) Sequence View

Sequences are displayed above the alignment location indicator. The dynamic yellow bar associated with the latter represents the region of the alignment that is currently visible. The green dot on the right hand side indicates what proportion of the alignment is visible. The consensus sequence of the alignment is displayed along the top of the sequence area, and directly under this the “+” indicates columns where all characters agree with the consensus character. Sequences and their titles are selectable and when a sequence is chosen it will be traced through the corresponding network as a yellow line. Sequences that pass through a user-selected node on the network are displayed with a red dot next to the title. Basic interactive features associated with the sequence display include the masking of characters that are the same as consensus, altering font size and altering space allocated to displaying titles; these are achieved through the “Navigation and Control” panel. Site locations are highlighted in yellow along the top of the interface.

#### (2) Network View

The network depicting sequence diversity within the alignment is displayed directly below the alignment location indicator. The associated orange bar of the latter represents the region of the alignment that is currently represented by the network. The region begins from the current sequence view and extends to the right-hand side in a manner that is dependent on the number of consecutive windows, as well as their width (figure 2, step i); windows being regions from which nodes reflecting diversity are created. Both these parameters can be interactively altered by the user. Window locations are highlighted in orange along the bottom of the interface.

**Figure 2:**
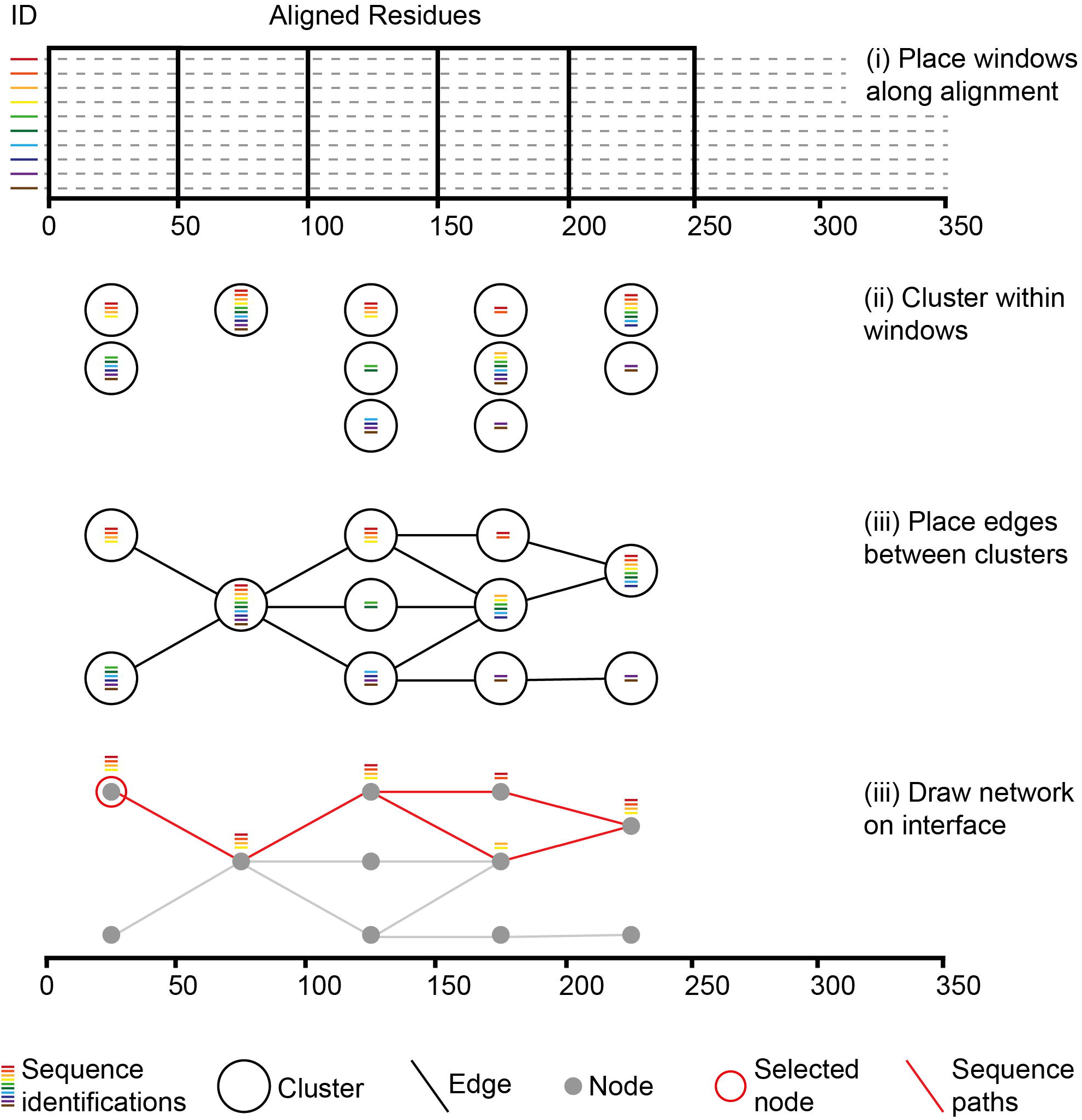
Network construction. Coloured bars indicate unique sequence id’s relative to the corresponding sequences (dotted lines). Within each window sequence id’s are associated with individual sequence fragments spanning that window (i) and fragments within windows are clustered (ii). Edges are placed between neighbouring clusters where they share one or more sequence id (iii). Clusters are represented visually on the network by grey dots. If a single cluster is selected the paths of all sequences passing through in relation to all other clusters (red lines) can be traced (vi).

##### (2.1) Nodes

For a given window clustering fragments of sequences that span it creates nodes (figure 2, step ii). Each cluster is created using an iterative approach. Initially a fragment is randomly selected to be a seed for a newly created empty cluster. All related fragments to that seed are then added to the cluster and become seeds for the next iteration. The metric used to define relatedness is hamming distance, in which the number of different characters between two aligned sequences are counted. The default threshold value is 0.3, indicating that fragments that have less than 30% divergence from a seed are included within the cluster. More advanced measures of genetic distance exist that account for proposed models of sequence evolution at both nucleotide and amino acid levels [24,25], but for the rapid clustering across windows placed along an alignment for the purpose of visualization hamming distance works well [26]. Iterations continue until no more next-round seeds can be identified. If unclustered fragments within the window still exist, a new cluster is initiated by selecting another random seed from the remaining fragments and the process is repeated.

For windows where six or less clusters are created, all clusters are displayed as grey coloured circles, or nodes, on the network. For windows with more than six clusters, the largest six are displayed as nodes whilst the remaining are placed into a holding structure that is used for visualization purpose only and that is displayed as a black circle. Internally all clusters contained within the latter are treated separately, for example in relation to tracking and highlighting paths. Here six was chosen to be the upper limit so that following edge placement (next section), and during edge crossover minimization, the maximum number of nodes that need order re-arrangement within any one window is seven, including the holding node. This is because for a set containing n items, there are n factorial different order permutations [27], and during edge crossover minimization the number of edge crossovers produced by each permutation, in relation to nodes within a neighbouring window, must be counted. For a given window if there are the maximum of seven nodes present, 7! permutations (5040) must be identified during crossover minimization and this can be done in a reasonable time (< 1 second on an average laptop). If on the other hand there are fifteen nodes allowed within a window, then there are 15! permutations (1307674368000) requiring a time of many days.

The default number (10) and width (50 bp) of windows, as well as the pairwise distance threshold, can be altered using the slider bars within the “Navigation and Control” area. The CViews graphical display is for rapid intuitive visualization, and if a higher clustering resolution is required, i.e. less than the 0.2 lower bound allowed for the display network, the user can perform this using the “Cluster Sequences” option of the “Alignment” menu. The number of sequence passing through each node is indicated in green on the left and right hand sides of the network.

##### (2.2) Edges

Edges are placed between nodes of neighbouring windows where they possess fragments that are derived from the same underlying sequence(s) (figure 2, iii). Consequently, individual sequences can be traced through nodes across different windows. As previously mentioned edge crossover between nodes within adjacent windows is minimized. Starting at the second most right-hand-side window, this is done by calculating all possible node order permutations, following which for each permutation, the number of edge crossovers to nodes within the adjacent right-hand window is counted, node layout order in the latter being kept constant (figure 3). The permutations that produce the minimum number of crossovers are selected (figure 3, red numbers), and from these a random one is used. The process is then repeated one window to the left, until the first window of the alignment is reached. Crossover minimization, while visually more pleasing, has no effect on the sequence information or underlying node connections. Following the connection of edges it is possible to click on nodes within the graph and track sequences that pass through them (figure 2, iv). On the interface, such sequence paths are displayed in red, and within the sequence display area a red dot are placed next to the titles of included sequences.

**Figure 3:**
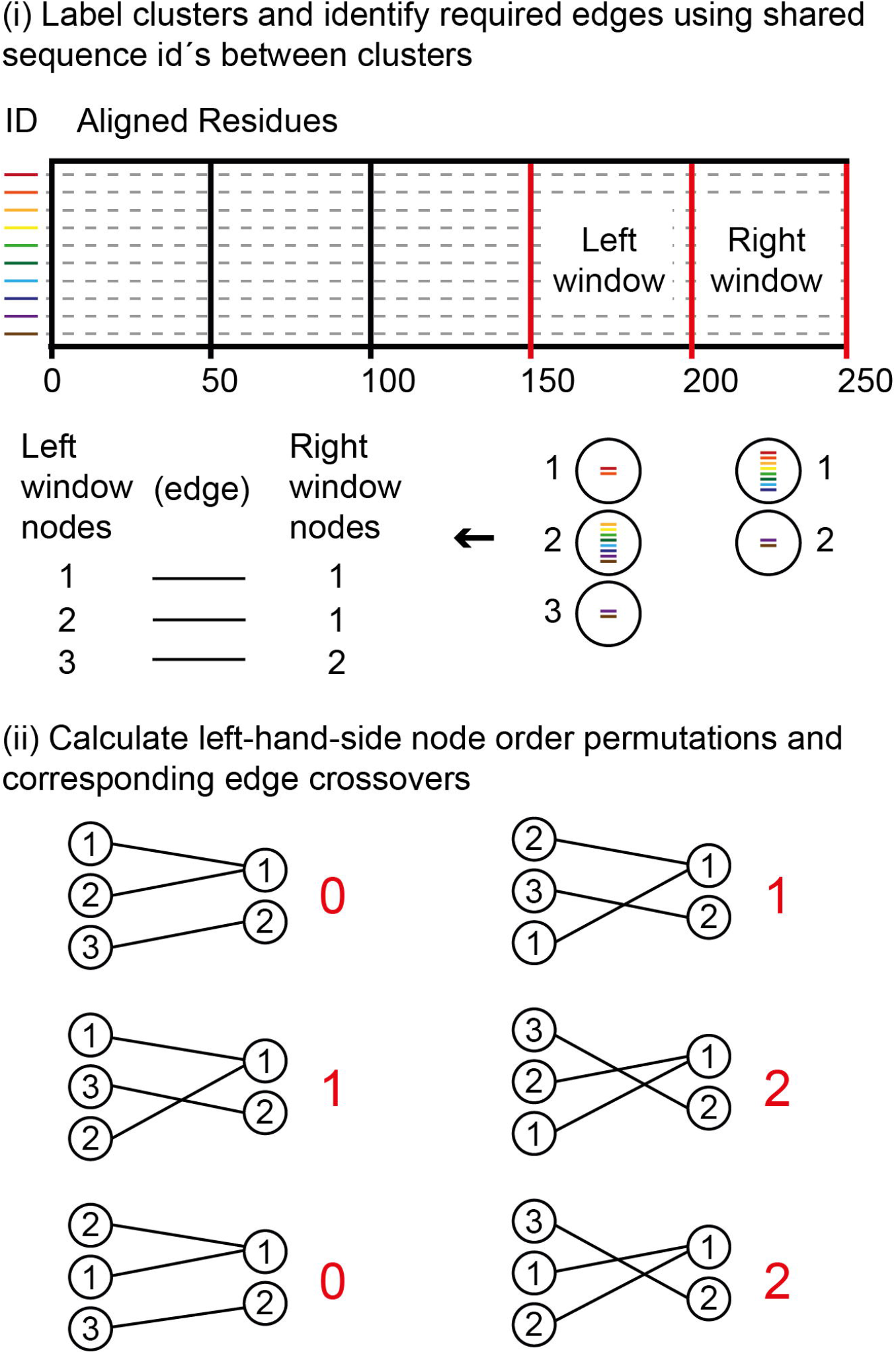
Minimization of edge crossovers between nodes of the two right most windows of the alignment. This process is repeated until the left most window (anchored on site 1) is reached. Clusters within the two windows are labelled with integers and required edges, based on the sequence ids (coloured bars), are listed (i). All order permutations of the current left window are identified and for each permutation the required edges are placed relative to the constant cluster order of the right window (ii). Crossovers are then counted (red numbers). Of the permutations that produce the minimum number of crossovers a random one is selected for graphical node layout order.

#### (3) Navigation and Control

Access to all the previous described options is provided through the slider bars associated with the navigation and control panel (bottom right of figure 1). The buttons labelled with then red directional arrows are used to scroll through the alignment. These were implemented to remove the need for flat scroll bars as future developments will be aimed at tablets and mobile devices. The red dot, at the centre of the four scroll arrows, immediately jumps a viewpoint at the centre of the alignment. In addition to the directional arrows the user can click directly on the grey squares along the alignment location indicator bar to immediately move to a particular location. Within this control area there are also three buttons used for printing the network to a .png formatted image file.

#### (4) Menu Driven Output

Output options are accessed through the top menu bar and can be applied to (i) the alignment in general, (ii) subsets of sequences whose titles match a user search tag, (iii) subsets of sequences that pass through a selected node and (iv) subsets of sequences defined by the user based on a supplied file of titles. In addition to exporting subsets of sequences and/or specified regions of the alignment Cview can generate summary statistics such as frequencies of residues and kmers and tertiary, pairwise distance matrix’s, variant count information and clustered sets of sequences. A detailed description of each output option is presented on the wiki associated with the SourceForge project page (https://sourceforge.net/p/cview/wiki/Help/).

## Results

### (1) The software

CView has been implemented in Java and runs on operating systems with installed Java Runtime Environment 8.0 or higher. It has been developed using an object-orientated approach for ease of plug-in development; where plug-ins related to alignment visualization will be based on user feedback. To obtain an executable jar file, download the cview.zip file from the SourceForge project page. Following the extraction the CView.jar file from the zip file, CView is executed by double clicking on the jar file. This will launch the interface through which alignments can be loaded. Alignments must be in fasta format and are be loaded using the “Load (fasta)” option of the “All Sequences” menu. Once a fasta-formatted alignment is loaded the workflow is driven by how the user interacts with the interface and the various output options. Test data, in the form of an alignment consisting of 636 sequences representing the gp120 region of the HIV-1 genome is included with the cview.zip file that contains the software. This data was obtained from the Los Alamos HIV sequence database [21].

### (2) Test Case Example: Exploring variation associated with co-receptor usage

#### (2.1) Background

HIV-1 viruses can be characterized into two phenotypes that are dependent on cellular tropism and that are as a result of differences in co-receptor usage [28]. The macrophage tropic phenotype, often referred to as R5, requires the CCR5 co-receptor, whilst the T-cell tropic phenotype (X4) uses the CXCR4 co-receptor, the latter often emerging later on during infection [29]. Co-receptor usage can be detected by computational analysis based on specific genetic alterations within the V3 loop of the gp120 gene [15,16]. Genetic variation within this region, of approximately 105 nt in length, lead to structural shifts that result in optimized binding to one co-receptor or the other [30]. For demonstrating the applicability of CView we have used it to explore and summarize this known variation relating to co-receptor usage from an alignment of HIV-1 subtype B gp120 sequences.

#### (2.2) Method

1. All North American subtype B gp120 sequences, verified to be CCR5-using sequences (n = 636), were downloaded from the Los Alamos HIV sequence database in aligned fasta format [21]. These were loaded into CView, using the “Load (fasta)” option of the “All Sequences” menu, following which they were saved in unaligned format using the “Save (unaligned)” option of the “All Sequences” menu. Additionally, the titles of these sequences were saved to a separate file using the “Save (titles)” option.
2. Step 1 was repeated for CXCR4-using sequences (n = 76).
3. In order to make sites directly comparable between the two sets of unaligned sequences, they were combined into a single file and aligned using MUSCLE [1].
4. The resulting alignment was loaded into CView and the consensus sequence of the region spanning the V3 loop was saved using the “Save (consensus)” option of the “All Sequences” menu. Within this alignment the co-ordinates of the region spanning the V3 loop were from 1436 to 1568. Although the exact location of the V3 loop within the gp120 region is known, the coordinates will vary depending on the alignment due to the placement of gaps during the alignment process. The exact co-ordinates for our alignment were identified by eye using the V3 sequence of the HIV-1 reference strain (Name: HXB2-LAI-IIIB-BRU, Accession: K03455), where the start residues of the loop are TGTACAAGACCC and the end residues are CAAGCACATTGT [21].
5. Using the original sequence titles saved in step 1, the proportion of the alignment corresponding to R5 sequences was saved in aligned format. This was done using the “Save (from - to)” option under the “Groups” menu item, where the titles were supplied as a list to define the group.
6. Step 4 was repeated for the titles corresponding to the X4 sequences.
7. Steps 5 and 6 resulted in two sub-alignments whose site co-ordinates are compatible as they were both extracted from the same underlying source. For each of the extracted R5 and X4 alignments, CView was used to obtain a list of all variants spanning the V3 loop along with their frequencies. This was done using the “Frequencies (variants)” option of the “All Sequences” menu, where the co-ordinates used were those described in step 3.
8. Variants were translated using the EMBOSS Transeq tool [31].
9. For each of the R5 and X4 alignments, nucleotide frequencies were obtained using the “Frequencies (nuc/aa)” option of the “All Sequence” menu (co-ordinates: 1436 to 1568).

The underlying alignment described for this use-case scenario, consisting of all the HIV-1 SUBTYPE B sequences spanning the GP120 region of the genome that have been verified as either being a CCR5-using (n = 636) or CXCR-using (n = 76), is available from the CView project page, within the compressed folder USE_CASE_DATA.zip. A further test dataset consisting of just the CCR5-using sequences from above is contained within the zip folder where the software itself is located.

#### (2.3) Result and Discussion

Figure 4A displays the consensus sequence of the V3 region from the MUSCLE generated alignment prior to being divided by genotype. The top ten most frequent variants from each of the two genotypes are also displayed. The seqPublish tool, located at https://www.hiv.lanl.gov/content/sequence/SeqPublish/seqpublish.html [21], was used to format these alignments from the CView output such that characters identical to those of the consensus sequence were hidden. A similar feature is available at the bottom of the output file that is generated by the “Variant Frequency” option of CView, where residues that are identical to those present on the most frequent variant are represented by a “|” character. The translation of each of these variants is presented within figure 4B. A summary, using sequence logos [32], is available within figure 4C where it can be observed that at translated site 11 the positively charged amino acid residues R (arginine) and H (histidine) are present within the sequences that were known to be CXCR4-using, while they are absent within the sequences obtained from the CCR5-uisng strains. At site 26 a similar observation is made in relation to positively charged residues, this time including a K (lysine) residue; although there is a minority K also present at 26 within the CCR5-using variants. This is a known observation where the presence of positively charged amino acids at sites 11 and 26 result in a structural alteration that optimizes CXCR4 co-receptor binding [15,16]. The steps leading to this observation, within this use-case scenario, demonstrate the utility of CView when exploring such alignment data. Complete per-site nucleotide frequencies for both R5 and X4 sequences spanning the V3 region are presented in supplementary table S1.

**Figure 4:**
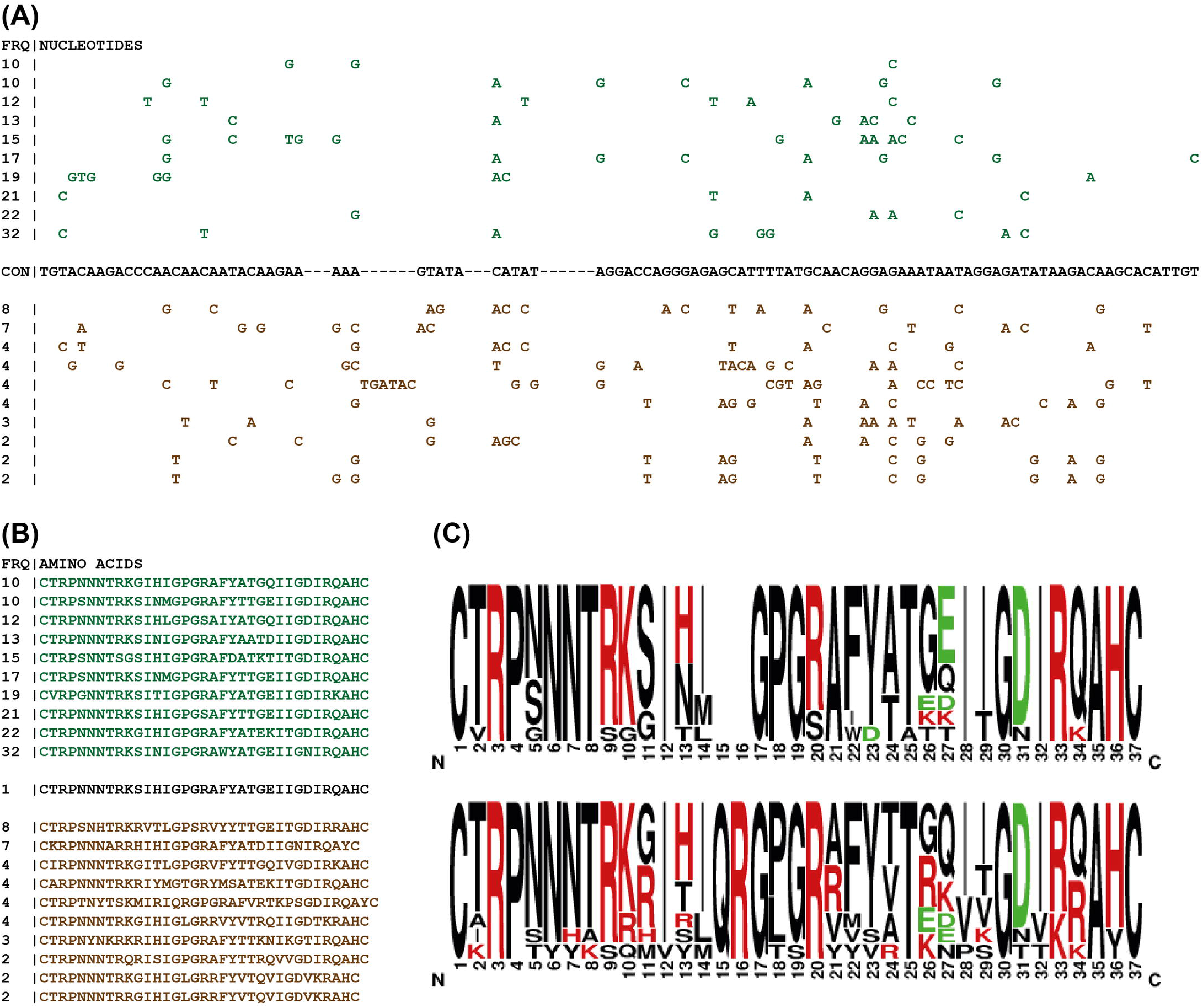
Summarization of variation present within the V3 loop. (A) Green residues represent non-consensus residues from the ten most frequent variants associate with the CCR5-using phenotype. Brown represents those of the CXCR4-using phenotype. The consensus sequence (black) is shown. (B) Translations of the most frequent ten variants from each phenotype. (C) Sequence logos summarizing these translations. The top logo is from represents the CCR5-using sequences whilst the bottom represents the CXCR4-using ones.

## Conclusion

CView is a tool that allows the user to interactively explore sequence alignments with the aid of a dynamic network that summarizes the diversity present. Here we have described how CView was designed and, as an example, we have used it to aid in the characterization of known variation between sequences involved in HIV-1 co-receptor usage. The exact usage scenario in which CView can be applied is dependent on the requirements of the individual user. CView is available from https://sourceforge.net/p/cview.

## Supporting Information Captions

**Table S1: Nucleotide frequencies from the V3 loop.** Sites covering codon 11 and 26 are highlighted in red.

