## Supplemental Table 1 for "CView: A network based tool for enhanced alignment visualization"

**Tabe S1: Nucleotide frequencies across the V3 loop.**

|  | | **CCR5 - using** | | | | | **CXCR4 - using** | | | | |
| --- | --- | --- | --- | --- | --- | --- | --- | --- | --- | --- | --- |
| **Site** | **codon** | **-** | **A** | **C** | **G** | **T** | **-** | **A** | **C** | **G** | **T** |
| 1436 | 1 | 1 | 0 | 0 | 0 | 634 | 0 | 0 | 0 | 0 | 75 |
| 1437 |  | 1 | 0 | 0 | 634 | 0 | 0 | 0 | 0 | 75 | 0 |
| 1438 |  | 1 | 0 | 69 | 0 | 564 | 0 | 0 | 5 | 0 | 70 |
| 1439 | 2 | 1 | 560 | 0 | 72 | 2 | 0 | 68 | 0 | 7 | 0 |
| 1440 |  | 1 | 14 | 520 | 0 | 100 | 0 | 12 | 55 | 0 | 8 |
| 1441 |  | 1 | 600 | 9 | 25 | 0 | 0 | 75 | 0 | 0 | 0 |
| 1442 | 3 | 1 | 633 | 1 | 0 | 0 | 0 | 75 | 0 | 0 | 0 |
| 1443 |  | 1 | 2 | 0 | 632 | 0 | 0 | 0 | 0 | 75 | 0 |
| 1444 |  | 1 | 576 | 0 | 58 | 0 | 0 | 67 | 0 | 8 | 0 |
| 1445 | 4 | 1 | 0 | 610 | 0 | 24 | 0 | 0 | 74 | 1 | 0 |
| 1446 |  | 1 | 4 | 604 | 0 | 26 | 0 | 1 | 73 | 0 | 1 |
| 1447 |  | 1 | 4 | 579 | 0 | 51 | 1 | 0 | 74 | 0 | 0 |
| 1448 | 5 | 1 | 570 | 20 | 39 | 5 | 1 | 67 | 0 | 6 | 1 |
| 1449 |  | 1 | 524 | 1 | 109 | 0 | 1 | 51 | 5 | 18 | 0 |
| 1450 |  | 1 | 0 | 588 | 0 | 46 | 0 | 0 | 70 | 0 | 5 |
| 1451 | 6 | 1 | 634 | 0 | 0 | 0 | 0 | 68 | 1 | 0 | 6 |
| 1452 |  | 1 | 633 | 0 | 0 | 1 | 0 | 74 | 0 | 1 | 0 |
| 1453 |  | 1 | 0 | 554 | 0 | 80 | 0 | 3 | 72 | 0 | 0 |
| 1454 | 7 | 1 | 633 | 0 | 0 | 1 | 0 | 59 | 10 | 1 | 5 |
| 1455 |  | 1 | 633 | 0 | 1 | 0 | 0 | 72 | 0 | 2 | 1 |
| 1456 |  | 1 | 0 | 53 | 1 | 580 | 0 | 0 | 7 | 1 | 67 |
| 1457 | 8 | 1 | 633 | 0 | 1 | 0 | 0 | 64 | 0 | 9 | 2 |
| 1458 |  | 1 | 1 | 632 | 0 | 1 | 0 | 5 | 62 | 0 | 8 |
| 1459 |  | 1 | 631 | 0 | 3 | 0 | 0 | 63 | 0 | 12 | 0 |
| 1460 | 9 | 1 | 626 | 8 | 0 | 0 | 0 | 75 | 0 | 0 | 0 |
| 1461 |  | 1 | 0 | 0 | 631 | 3 | 0 | 2 | 1 | 72 | 0 |
| 1462 |  | 1 | 528 | 5 | 76 | 25 | 0 | 62 | 9 | 3 | 1 |
| 1463 | 10 | 1 | 613 | 4 | 17 | 0 | 0 | 71 | 4 | 0 | 0 |
| 1464 |  | 634 | 1 | 0 | 0 | 0 | 72 | 2 | 0 | 0 | 1 |
| 1465 |  | 634 | 1 | 0 | 0 | 0 | 72 | 3 | 0 | 0 | 0 |
| 1466 | 11 | 634 | 1 | 0 | 0 | 0 | 72 | 2 | 1 | 0 | 0 |
| 1467 |  | 0 | 561 | 12 | 62 | 0 | 0 | 56 | 0 | 19 | 0 |
| 1468 |  | 0 | 602 | 1 | 32 | 0 | 0 | 68 | 1 | 6 | 0 |
| 1469 | 12 | 0 | 474 | 2 | 159 | 0 | 0 | 38 | 18 | 19 | 0 |
| 1470 |  | 635 | 0 | 0 | 0 | 0 | 70 | 0 | 0 | 0 | 5 |
| 1471 |  | 635 | 0 | 0 | 0 | 0 | 70 | 0 | 0 | 5 | 0 |
| 1472 | 13 | 635 | 0 | 0 | 0 | 0 | 70 | 5 | 0 | 0 | 0 |
| 1473 |  | 635 | 0 | 0 | 0 | 0 | 70 | 0 | 0 | 0 | 5 |
| 1474 |  | 635 | 0 | 0 | 0 | 0 | 70 | 5 | 0 | 0 | 0 |
| 1475 | 14 | 635 | 0 | 0 | 0 | 0 | 70 | 0 | 5 | 0 | 0 |
| 1476 |  | 0 | 0 | 0 | 634 | 1 | 0 | 11 | 0 | 62 | 2 |
| 1477 |  | 0 | 2 | 44 | 2 | 587 | 0 | 13 | 12 | 17 | 33 |
| 1478 | 15 | 0 | 629 | 0 | 5 | 1 | 0 | 64 | 0 | 11 | 0 |
| 1479 |  | 0 | 0 | 0 | 0 | 635 | 0 | 0 | 0 | 0 | 75 |
| 1480 |  | 0 | 617 | 17 | 1 | 0 | 0 | 74 | 0 | 1 | 0 |
| 1481 | 16 | 634 | 1 | 0 | 0 | 0 | 74 | 1 | 0 | 0 | 0 |
| 1482 |  | 635 | 0 | 0 | 0 | 0 | 74 | 0 | 0 | 0 | 1 |
| 1483 |  | 635 | 0 | 0 | 0 | 0 | 74 | 0 | 0 | 1 | 0 |
| 1484 | 17 | 0 | 166 | 460 | 2 | 7 | 0 | 24 | 41 | 0 | 10 |
| 1485 |  | 0 | 541 | 80 | 14 | 0 | 0 | 47 | 20 | 7 | 1 |
| 1486 |  | 0 | 1 | 14 | 11 | 609 | 0 | 1 | 7 | 5 | 62 |
| 1487 | 18 | 0 | 584 | 16 | 4 | 31 | 0 | 59 | 14 | 2 | 0 |
| 1488 |  | 0 | 0 | 1 | 0 | 634 | 0 | 0 | 1 | 6 | 68 |
| 1489 |  | 629 | 6 | 0 | 0 | 0 | 75 | 0 | 0 | 0 | 0 |
| 1490 | 19 | 629 | 0 | 0 | 6 | 0 | 75 | 0 | 0 | 0 | 0 |
| 1491 |  | 629 | 0 | 0 | 6 | 0 | 75 | 0 | 0 | 0 | 0 |
| 1492 |  | 629 | 6 | 0 | 0 | 0 | 75 | 0 | 0 | 0 | 0 |
| 1493 | 20 | 629 | 0 | 6 | 0 | 0 | 75 | 0 | 0 | 0 | 0 |
| 1494 |  | 629 | 0 | 6 | 0 | 0 | 75 | 0 | 0 | 0 | 0 |
| 1495 |  | 1 | 503 | 2 | 127 | 2 | 0 | 60 | 0 | 15 | 0 |
| 1496 | 21 | 0 | 0 | 6 | 629 | 0 | 0 | 0 | 0 | 75 | 0 |
| 1497 |  | 0 | 7 | 20 | 608 | 0 | 0 | 0 | 0 | 75 | 0 |
| 1498 |  | 0 | 627 | 0 | 8 | 0 | 0 | 74 | 0 | 1 | 0 |
| 1499 | 22 | 0 | 0 | 630 | 2 | 3 | 0 | 7 | 68 | 0 | 0 |
| 1500 |  | 0 | 2 | 628 | 2 | 3 | 0 | 0 | 64 | 0 | 11 |
| 1501 |  | 0 | 613 | 1 | 21 | 0 | 0 | 74 | 0 | 1 | 0 |
| 1502 | 23 | 0 | 0 | 0 | 635 | 0 | 0 | 11 | 0 | 64 | 0 |
| 1503 |  | 0 | 0 | 0 | 635 | 0 | 0 | 0 | 0 | 75 | 0 |
| 1504 |  | 0 | 28 | 59 | 545 | 3 | 0 | 1 | 11 | 63 | 0 |
| 1505 | 24 | 0 | 602 | 3 | 30 | 0 | 0 | 74 | 1 | 0 | 0 |
| 1506 |  | 0 | 56 | 2 | 577 | 0 | 0 | 0 | 0 | 75 | 0 |
| 1507 |  | 1 | 483 | 31 | 65 | 55 | 1 | 73 | 0 | 1 | 0 |
| 1508 | 25 | 1 | 90 | 1 | 542 | 1 | 1 | 12 | 1 | 55 | 6 |
| 1509 |  | 1 | 1 | 611 | 1 | 21 | 1 | 6 | 41 | 11 | 16 |
| 1510 |  | 0 | 627 | 6 | 2 | 0 | 0 | 69 | 6 | 0 | 0 |
| 1511 | 26 | 0 | 46 | 7 | 0 | 582 | 0 | 8 | 1 | 5 | 61 |
| 1512 |  | 0 | 3 | 0 | 57 | 575 | 0 | 11 | 0 | 0 | 64 |
| 1513 |  | 2 | 11 | 11 | 98 | 513 | 0 | 0 | 6 | 7 | 62 |
| 1514 | 27 | 1 | 0 | 4 | 25 | 605 | 0 | 0 | 5 | 7 | 63 |
| 1515 |  | 1 | 619 | 1 | 1 | 13 | 0 | 57 | 6 | 4 | 8 |
| 1516 |  | 0 | 0 | 19 | 1 | 615 | 0 | 0 | 0 | 0 | 75 |
| 1517 | 28 | 0 | 159 | 0 | 469 | 7 | 0 | 36 | 0 | 38 | 1 |
| 1518 |  | 0 | 0 | 627 | 0 | 8 | 0 | 0 | 59 | 5 | 11 |
| 1519 |  | 3 | 629 | 1 | 1 | 1 | 0 | 63 | 12 | 0 | 0 |
| 1520 | 29 | 3 | 589 | 0 | 42 | 1 | 0 | 74 | 0 | 0 | 1 |
| 1521 |  | 3 | 0 | 629 | 2 | 1 | 0 | 0 | 75 | 0 | 0 |
| 1522 |  | 634 | 1 | 0 | 0 | 0 | 75 | 0 | 0 | 0 | 0 |
| 1523 | 30 | 634 | 1 | 0 | 0 | 0 | 75 | 0 | 0 | 0 | 0 |
| 1524 |  | 634 | 0 | 0 | 1 | 0 | 75 | 0 | 0 | 0 | 0 |
| 1525 |  | 634 | 1 | 0 | 0 | 0 | 75 | 0 | 0 | 0 | 0 |
| 1526 | 31 | 634 | 1 | 0 | 0 | 0 | 75 | 0 | 0 | 0 | 0 |
| 1527 |  | 634 | 0 | 1 | 0 | 0 | 75 | 0 | 0 | 0 | 0 |
| 1528 |  | 634 | 1 | 0 | 0 | 0 | 75 | 0 | 0 | 0 | 0 |
| 1529 | 32 | 634 | 1 | 0 | 0 | 0 | 75 | 0 | 0 | 0 | 0 |
| 1530 |  | 634 | 0 | 0 | 1 | 0 | 75 | 0 | 0 | 0 | 0 |
| 1531 |  | 39 | 596 | 0 | 0 | 0 | 28 | 47 | 0 | 0 | 0 |
| 1532 | 33 | 39 | 50 | 0 | 546 | 0 | 28 | 19 | 0 | 28 | 0 |
| 1533 |  | 39 | 65 | 25 | 506 | 0 | 28 | 14 | 0 | 33 | 0 |
| 1534 |  | 0 | 544 | 3 | 86 | 2 | 0 | 64 | 1 | 10 | 0 |
| 1535 | 34 | 0 | 87 | 83 | 465 | 0 | 0 | 21 | 25 | 29 | 0 |
| 1536 |  | 0 | 566 | 39 | 30 | 0 | 0 | 71 | 0 | 4 | 0 |
| 1537 |  | 0 | 446 | 146 | 7 | 36 | 0 | 54 | 4 | 0 | 17 |
| 1538 | 35 | 0 | 595 | 0 | 40 | 0 | 0 | 55 | 5 | 15 | 0 |
| 1539 |  | 0 | 3 | 0 | 0 | 632 | 1 | 0 | 6 | 0 | 68 |
| 1540 |  | 0 | 592 | 43 | 0 | 0 | 3 | 72 | 0 | 0 | 0 |
| 1541 | 36 | 0 | 630 | 0 | 5 | 0 | 3 | 61 | 0 | 6 | 5 |
| 1542 |  | 0 | 0 | 57 | 1 | 577 | 3 | 3 | 23 | 1 | 45 |
| 1543 |  | 0 | 635 | 0 | 0 | 0 | 1 | 73 | 0 | 1 | 0 |
| 1544 | 37 | 0 | 0 | 0 | 635 | 0 | 1 | 0 | 0 | 74 | 0 |
| 1545 |  | 0 | 0 | 0 | 635 | 0 | 0 | 0 | 0 | 75 | 0 |
| 1546 |  | 0 | 553 | 0 | 81 | 1 | 0 | 73 | 0 | 2 | 0 |
| 1547 | 38 | 0 | 126 | 1 | 508 | 0 | 0 | 19 | 0 | 56 | 0 |
| 1548 |  | 0 | 635 | 0 | 0 | 0 | 0 | 71 | 4 | 0 | 0 |
| 1549 |  | 0 | 2 | 90 | 1 | 542 | 0 | 0 | 13 | 0 | 62 |
| 1550 | 39 | 0 | 634 | 0 | 1 | 0 | 0 | 69 | 0 | 6 | 0 |
| 1551 |  | 0 | 0 | 1 | 0 | 634 | 0 | 0 | 5 | 0 | 70 |
| 1552 |  | 0 | 633 | 0 | 2 | 0 | 0 | 74 | 0 | 1 | 0 |
| 1553 | 40 | 0 | 633 | 2 | 0 | 0 | 0 | 75 | 0 | 0 | 0 |
| 1554 |  | 0 | 3 | 0 | 632 | 0 | 0 | 11 | 0 | 63 | 1 |
| 1555 |  | 0 | 588 | 0 | 47 | 0 | 0 | 74 | 0 | 1 | 0 |
| 1556 | 41 | 0 | 80 | 555 | 0 | 0 | 0 | 12 | 63 | 0 | 0 |
| 1557 |  | 0 | 611 | 0 | 20 | 4 | 0 | 49 | 0 | 26 | 0 |
| 1558 |  | 0 | 605 | 0 | 30 | 0 | 0 | 69 | 1 | 5 | 0 |
| 1559 | 42 | 0 | 0 | 0 | 635 | 0 | 0 | 0 | 0 | 75 | 0 |
| 1560 |  | 0 | 0 | 634 | 0 | 1 | 0 | 0 | 75 | 0 | 0 |
| 1561 |  | 0 | 629 | 2 | 2 | 2 | 0 | 74 | 0 | 1 | 0 |
| 1562 | 43 | 0 | 2 | 551 | 0 | 82 | 0 | 0 | 56 | 0 | 19 |
| 1563 |  | 0 | 610 | 24 | 0 | 1 | 0 | 74 | 0 | 1 | 0 |
| 1564 |  | 0 | 0 | 15 | 0 | 620 | 0 | 0 | 0 | 1 | 74 |
| 1565 | 44 | 0 | 0 | 0 | 0 | 635 | 0 | 0 | 0 | 0 | 75 |
| 1566 |  | 0 | 0 | 0 | 635 | 0 | 0 | 0 | 0 | 75 | 0 |
| 1567 |  | 0 | 0 | 26 | 0 | 609 | 0 | 0 | 2 | 0 | 73 |
| 1568 | 45 | 0 | 609 | 0 | 26 | 0 | 0 | 75 | 0 | 0 | 0 |
